# SCOT+: A Comprehensive Software Suite for Single-Cell alignment Using Optimal Transport

**DOI:** 10.1101/2025.05.21.655322

**Authors:** Colin Baker, Tuan Pham, Pinar Demetci, Quang Huy Tran, Ievgen Redko, Bjorn Sandstede, Ritambhara Singh

## Abstract

**Summary:** New advances in single-cell multi-omics experiments have allowed biologists to examine how various biological factors regulate processes in concert on the cellular level. However, measuring multiple cellular features for a single cell can be quite resource-intensive or impossible with the current technology. By using optimal transport (OT) to align cells and features across disparate datasets produced by separate assays, Single Cell alignment using Optimal Transport+ (SCOT+), our unsupervised single-cell alignment software suite, allows biologists to align their data without the need for any correspondence. SCOT+ has a generic optimal transport solution that can be reduced to multiple different OT optimization procedures, each of which provide state-of-the-art single-cell alignment performance. With our user-friendly website and tutorials, this new package will help improve biological analyses by allowing for more accurate downstream analyses on multi-omics single-cell measurements.

**Implementation and Availability:** Our algorithm is implemented in Pytorch and available on PyPI and GitHub (https://github.com/scotplus/scotplus). Additionally, we have many tutorials available in a separate GitHub repository (https://github.com/scotplus/book_source) and on our website (https://scotplus.github.io/).

## Introduction

Single-cell multi-omic study integrates various modalities of omics, such as genomic, transcriptomic, proteomic, or epigenomic data, at the single-cell resolution [1, 2, 3]. Such an integration is crucial for biologists to gain a deeper understanding of cell regulation and development. However, simultaneously measuring multi-modal data for single cells is either difficult or expensive due to technical limitations [1]. Thus researchers rely on separately sequenced single-cell measurements that have to be computationally integrated.

As a result, several computational methods have been developed to integrate multiple single-cell datasets that draw from similar cell populations [4, 5, 6, 7]. A single-cell dataset is usually represented as a count matrix where the “sample” (row) is a specific cell and the “feature” (column) is a genomic measurement on this cell. To integrate two single-cell datasets, existing methods reconstruct a unified matrix that combines samples and feature sets from both. This requires the assumption that there exist some meaningful relationships between each set of features. However, this is a difficult problem because such relationships are not observable across disparate sample sets without 1-1 correspondence of cells. In order to combat such issues, supervised methods like Seurat [4] and Harmony [5] utilize explicit, ground-truth sample or feature correspondences between the two datasets to extrapolate the rest of the alignment. On the other hand, unsupervised methods like MMD-MA [6] and BindSC [7] attempt to estimate a multi-model dataset without any prior sample or feature correspondence information.

Among the existing unsupervised single-cell multi-omic alignment methods, optimal transport (OT) approaches [8, 9, 10, 11, 12, 13] have shown state-of-the-art performance for aligning separately profiled multi-omic datasets. Optimal transport constructs a probabilistic correspondence matrix between the data points of two input domains in order to match them in the most cost effective way possible [14]. A popular view of the problem is to imagine moving a pile of sand to fill in a hole through the least amount of work.

In the single-cell context, OT-based approaches allow for both distribution-free alignments and additional probabilistic interpretation of the coupling matrix prior to integration. Prior methods like UnionCom [8], Pamona [9], and uniPort [15] also leverage these properties to compute single-cell alignments. These methods have proven useful to, for example, co-embed and compare single-cell datasets across different conditions [16]. However, many of these different OT methods for alignment are scattered throughout different packages. The lack of a single, easy to use package that unifies such methods has made OT less accessible to the single-cell community.

Given the advantages of OT-based methods, we introduce **S**ingle **C**ell alignment using **O**ptimal **T**ransport+ (or SCOT+), a software suite that leverages three different OT formulations to integrate single-cell datasets. (1) Gromov-Wasserstein (GW) OT, applied for the single-cell integration task as SCOT [10], is best used in contexts where the geometric structure of the two datasets is important, while potential feature relationships between the two domains to be aligned could be nonlinear and but are not as relevant to alignment [17, 18, 19]. (2) Co-Optimal Transport (COOT), applied as SCOOTR [12], is best used in contexts where potential feature relationships are close to linear and well-known so that linear supervision might aid alignment more meaningfully [20]. Finally, (3) Augmented Gromov-Wasserstein (AGW) is a convex combination of GW and COOT distance that allows for feature supervision without any restriction to linearity and therefore brings together the benefits of both formulations at the cost of an extra hyperparameter [13]. Additionally, each of these formulations is applicable in cases where there is a disproportionate distribution of cell types in either dataset. Unbalancing a formulation, in this case, improves alignment, as shown in the SCOTv2 and UCOOT papers [11, 21, 22, 23]. The unbalanced version of AGW, which we term as UAGW, is a novel construct that arose from our generalization of the single-cell OT pipelines for SCOT+. It is the only single-cell OT formulation thus far that allows for unbalancing in the feature space. For example, it can account for cases where one gene might regulate a number of other genes at once resulting in a one-to-many correspondence.

SCOT+ provides a unified platform to access the best solutions to each of these formulations, so that users have many options available for integrating different types of single-cell datasets. Rather than having to handle significant hyperparameter tuning to appropriately align their data, with SCOT+, users can choose the formulation that more intuitively applies to their data for the best results. Additionally, SCOT+ utilizes a single, generic optimizer to solve all of its formulations, making it more user-friendly and accessible. We leverage our insight that all other OT formulations mentioned above are partial functions of UAGW, which we introduce in this paper and SCOT+ solves on the backend.

We show that SCOT+ performs well on real world single-cell datasets, both with respect to replicating co-assay data and correctly mapping cell type clusters across domains for separately sequenced single-cell measurements. We also discuss the potential for downstream analysis of the estimated multi-omic mappings for the community.

## Methods

### SCOT+ Optimal Transport Setup

Given *X* and *Y*, the count matrices from two single-cell measurements, SCOT+ seeks to learn a probabilistic pairing between their cells (and features) that integrates the two datasets. We assume below that *X* has *n*_*x*_ cells and *d*_*x*_ features, while *Y* has *n*_*y*_ cells and *d*_*y*_ features. Each of the GW, COOT, and AGW formulations in SCOT+ treat their input domains *X* and *Y* as probability distributions. By minimizing the cost of transporting cells from one domain to another, these methods produce a *coupling matrix* that is the learned pairing between them. We first examine the standard optimal transport problem [24, 18] that is defined as:

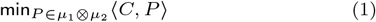

where *C* is some cost matrix such that *C*_*ij*_ is the cost of transporting cell *i* to cell *j*, and *μ*_1_, *μ*_2_ are the marginals for the solution to the problem. A coupling matrix *P*, which we force to have marginals *μ*_1_ and *μ*_2_, maps the cells across disparate sample spaces. Here, ⟨*C, P* ⟩ := ∑_*ij*_ *C*_*ij*_ *P*_*ij*_. This problem can also be generalized [25] as follows:

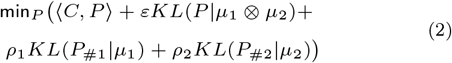

Here, the first Kullback–Leibler (KL) divergence term *KL*(*P* |*μ*_1_ ⊗ *μ*_2_) is for regularization. It controls the sparsity in *P*, making the problem more computationally tractable. Additionally, the *ρ*_1_*KL*(*P*_#1_|*μ*_1_) and *ρ*_2_*KL*(*P*_#2_|*μ*_2_) terms relax the constraints on *P*’s marginals. They allow flexibility of cell-cell mappings when there is a disproportionate cell-type representation across the datasets. Note that we use *P*_#1_ to represent the marginal distribution of *P* in the first domain and *P*_#2_ to represent the marginal distribution of *P* in the second domain. For example, *P*_#1_ = ∑_*j*_ *P*_*ij*_.

Another useful feature is that the user can provide supervision when learning the coupling matrix by adding the term *β*⟨*D, P* ⟩ to Equation (2). Here, *D* is some user-constructed matrix according to known cell-cell or feature-feature mappings and *β* is a coefficient for the magnitude of supervision [19].

Each of the SCOT+ formulations break down into solving the general problem presented in Equation (2) at each iteration. As a result, SCOT+ currently utilizes Sinkhorn’s algorithm [25] as its main engine. To retrieve an alignment from a coupling matrix *P* mapping *X* to *Y*, we take the weighted average of the genomic measurements in *Y* according to the weights in *P* [26, 27]. This produces a new matrix *Ŷ* with the *d*_*y*_ features of *Y* on the *n*_*x*_ cells of *X*:

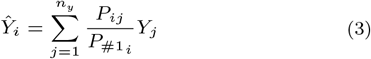

Where the index *i* refers to cells in *X* and *j* refers to the cells in *Y*. More information on this projection is presented in Supp. Section 3. In the next section, we take the general formulation designed for SCOT+ in Equation (2) and show how it can be easily extended to yield six different OT formulations relevant for different single-cell integration tasks.

#### Different SCOT+ Formulations

First, the GW OT formulation that SCOT+ utilizes to align count matrices *X* and *Y* is the following minimization problem [17, 28]:

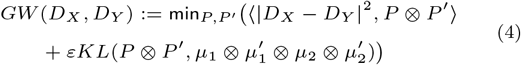

In this case, *D*_*X*_ and *D*_*Y*_ are intra-domain distance matrices that capture the local geometry of each domain. We calculate these by constructing a graph of cells in Euclidean space for each domain and computing the shortest path distances between two cells in the graph. SCOT+ aligns these two graphs by mapping their internal local geometries onto each other. This formulation encourages the minimization of the sum:

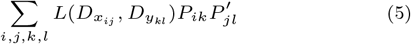

where *L* is the Euclidean norm, so that SCOT+ outputs an alignment of samples encoded by *P* and *P* ^*′*^ which minimizes the distortions between the local geometry in the domain *X* and the one in *Y*. Note that this formulation can also be marginally relaxed with the additional terms 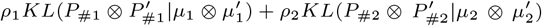 [21]. While GW and its unbalanced counterpart (UGW) are very effective for aligning cells, they do not explicitly produce any feature mappings. Such feature mappings, e.g. gene to chromatin regions, could be useful for biological hypothesis generation. Therefore, our next formulation addresses this gap.

SCOT+ also includes the COOT formulation [20] that learns two coupling matrices, one for cells and the other for features:

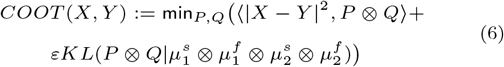

In this case, *Q* is a coupling matrix for the features of *X* and *Y* so that we minimize the joint transport of cells and features from one domain to another. We can also relax this formulation by adding 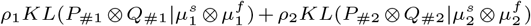 [23] resulting in its unbalanced form (or UCOOT). Unfortunately, COOT formulation can be sensitive to outliers and the magnitude of features in each domain [23].

Finally, the newest formulation utilized by SCOT+, AGW OT, merges these two loss functions (GW and COOT) as follows [13]:

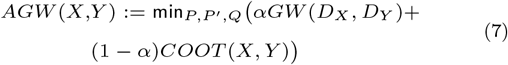

This formulation is more robust to outliers as the GW term preserves the local geometry and it also produces feature coupling matrices [13]. We introduce a novel relaxed AGW formulation by individually relaxing the GW and COOT terms as in the above descriptions. Our general optimization approach designed for SCOT+ lends itself to this novel unbalanced version of AGW (or UAGW). More details are provided in Supp. Section 1.

The availability of each of these formulations through a single SCOT+ package allows users to tailor their integration details to the specifics of their dataset. For example, one user with disproportionate cell types and a prior on the feature relationships might use UCOOT, but another with uniform cell types and no prior on the features might use GW.

## Results

We now demonstrate the versatility of SCOT+ for the integration of real-world single-cell datasets. We specifically focus on balanced and unbalanced scenarios where GW and AGW are applicable. For details on the hyperparameters we used, see Supp. Section 2.

### Cell-cell alignment on balanced datasets

As formalized above, SCOT+ GW OT aligns disparate single-cell measurements in a way that maintains the cell-cell similarities upon integration. In many cases where we do not need feature-feature relationships, GW can provide meaningful cell-cell alignments.

Specifically, balanced GW is applicable in cases where we have similar cell-type proportions across domains. To showcase this, we use a PBMC co-assay dataset [2] that simultaneously measures scRNA-seq and scATAC-seq data. This dataset has a 1-1 correspondence between single cells across these two domains, which we use only to quantify the alignment performance.

To execute an alignment between these domains, we split a subset of the PBMC dataset into two separate count matrices with the same set of 2407 cells. Then, we applied PCA to the scRNA-seq data to select the 50 most contributing components and topic modeling [29] to the scATAC-seq data to embed the domain into a smaller space with 50 topics. This reduces the dimension of the data to remove noise.

Next, we run SCOT+’s GW formulation and obtained a sample-level alignment across our two separate domains. A barycentric projection using the optimized coupling matrix returns a new single-cell dataset with the same set of original samples from the scATAC-seq data, but the features of the scRNA-seq data. We call this data the projected scATAC-seq data, which is visualized in Figure 1B.

**Fig. 1.**
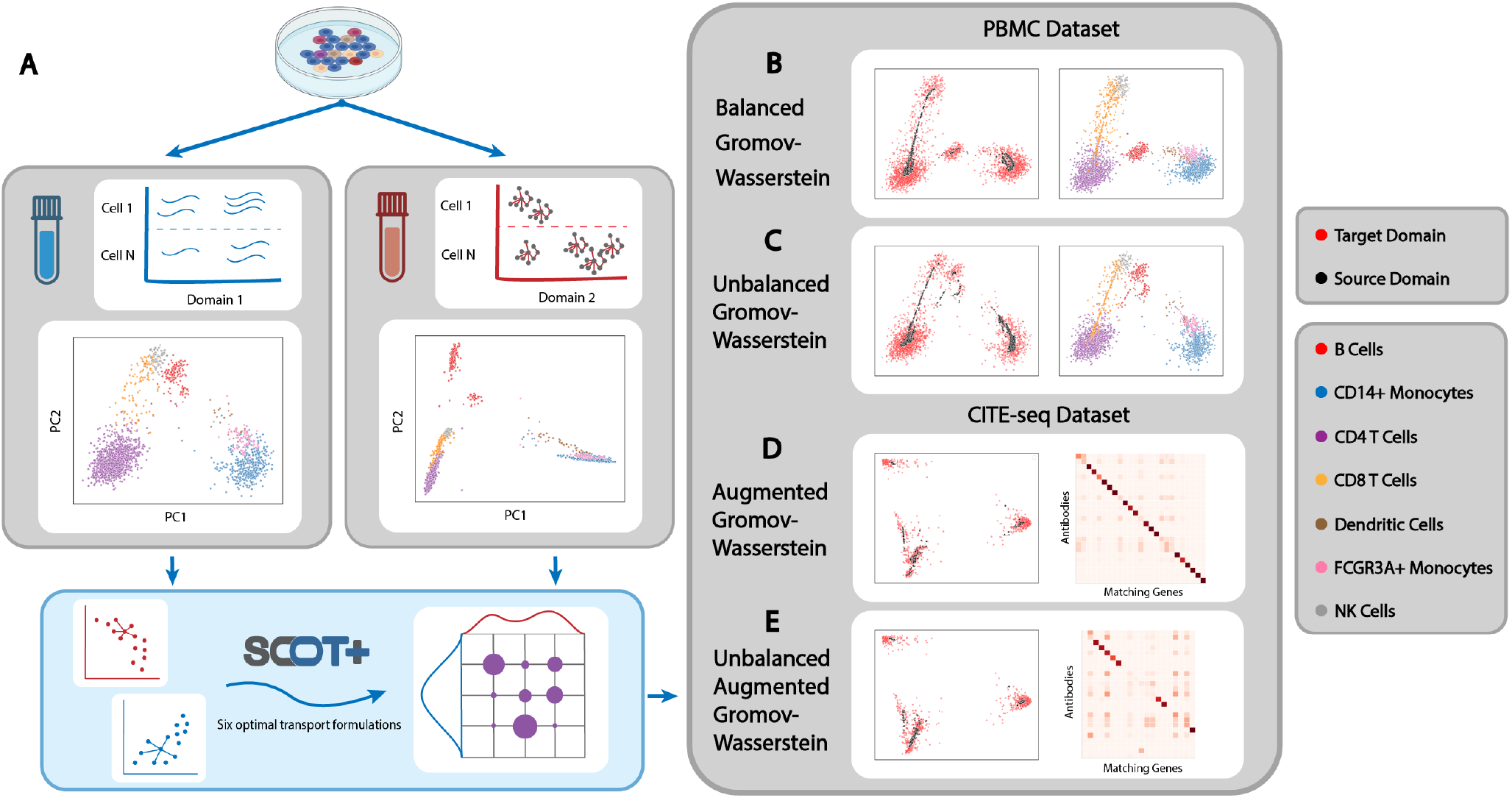
An example of the SCOT+ workflow on a PBMC dataset [2] and a CITE-seq dataset [3]. All plotted results highlight the projection of the data onto the top two principal components. **A** highlights the SCOT+ workflow. **B** displays the results of running GW on the PBMC data (scRNA-seq and scATAC-seq) in a balanced fashion, followed by a barycentric projection of the accessibility domain onto the gene expression domain. Note that the rightmost GW figure highlights the retained clustering of each cell type when we examine the full aligned dataset. GW reconstructs the gene expression domain from accessibility with low alignment error. **C** displays the results of running UGW on the same two domains after downsampling each cell type by a random amount. UGW reconstructs each cell type cluster with high accuracy. **D** displays the results of balanced AGW on the two CITE-seq domains (scRNA-seq and ADT), followed by a barycentric projection of the gene expression domain onto the ADT domain. AGW reconstructs the ADT domain from accessibility low alignment error and 73% of the feature matrix mass assigned to ground-truth feature relationships. **E** displays the results of running UAGW on the same two domains when we remove five ADT features. UAGW reconstructs the ADT domain with low alignment error and 52% feature mass assigned to ground-truth relationships.

To quantify the quality of this projected scATAC-seq data, we calculated the fraction of samples closer than the true match (FOSCTTM) using the ground-truth sample correspondence of the PBMC data. FOSCTTM measures the fraction of samples in the projected scATAC-seq data that are closer to a given sample in the original scRNA-seq data than its true match [10]. When this metric is lower, it means that each sample in the respective domains is relatively close to its correct match. In this example, we attained a FOSCTTM of 0.12, meaning that the projected scATAC-seq data had a similar geometry and shape to the original scRNA-seq data. This FOSCTTM value is comparable to the previous results in [13].

### Cell-cell alignment on unbalanced datasets

The balanced formulation of GW OT fails to account for disproportionate cell-type representation that usually exists in real-world datasets [11]. The unbalanced case of UGW (implemented in SCOT+) closely models what we expect biologists will observe under real-world conditions in separately sequenced single-cell datasets.

To simulate the unbalanced setting, we systematically downsampled both the scRNA-seq and scATAC-seq data from our previous example by cell type. In particular, we drew a random fraction between 0 and 0.5 for each cell type and discarded the corresponding number of samples in the scRNA-seq data. We then repeated the same process with different fractions for the scATAC-seq data and used UGW to compute the alignment and barycentric projections shown in Figure 1C.

In this case, we used a metric called label transfer accuracy (LTA) [8] to score our alignment as we do not have exact 1-1 correspondences anymore. To compute LTA, we trained a *k*-nn classifier on the original domain to be projected, where *k* is the actual number of cell types. Next, the score is determined by the accuracy of this classifier on the projected domain. In this example, SCOT+ achieved an LTA of 92.5%, meaning that it succeeds in transporting cell types from one domain to another in a more realistic single-cell integration setting.

### Feature-based alignment on balanced datasets

While GW and UGW work well in cases where we mostly care about obtaining a cell-cell alignment, AGW works better in cases where there are well-defined feature relationships [13]. In particular, AGW learns both cell-cell and feature-feature mappings, which is relevant for hypothesis generation in biology. Furthermore, this joint formulation allows for correspondence on either level to inform the other.

To show the usability of SCOT+’s AGW, we used a CITE-seq dataset [3], which has co-assayed scRNA-seq and ADT (antibody) data. We downsampled the features of both domains to 25 gene-antibody pairs that are known to be biologically related. After running AGW on these two datasets, we got a FOSCTTM of 0.08, but more importantly, we recovered the accurate feature map displayed in Figure 1D, which has 73% of its mass assigned to biologically validated antibody-gene pairs [13]. This example illustrates the capacity for SCOT+ to learn meaningful feature relationships whilst also integrating single domain datasets. It also demonstrates the additional cell-level alignment power gained from integrating feature information.

### Feature-based alignment on unbalanced datasets

AGW’s unbalanced counterpart, UAGW in SCOT+, is a new construct that arose from the generalization of our optimization framework. With the fully generic version of our solver, we can get impressive cell-cell alignments in the unbalanced feature setting.

To display this functionality, we downsampled the features of *Y* from our previous experiment to contain only 20 antibodies. After running UAGW on this data, where we balanced the GW term and left the COOT term unbalanced, we recovered a cell-level alignment as seen in Figure 1E. This alignment had FOSCTTM 0.09 and a reasonable feature coupling matrix, which had 52% of its mass assigned to known antibody-gene pairs [13]. If we had instead generated the coupling matrix randomly, the expected mass assigned to these known pairs would instead be 4%, indicating that UAGW assigned significant additional mass to these known relationships. In some settings without a known correspondence, cells with exaggerated mass relative to random could help generate hypotheses about feature relationships. As a result, we found that UAGW can successfully recover unbalanced feature relationships. UAGW additionally allows for supervision of the feature matrix in an unbalanced fashion, such that we can improve cell-level alignments.

## Discussion

With SCOT+, we believe that the single-cell community will be better able to understand and utilize the end-to-end alignment procedure of OT methods. Since it uses only one generic solution in the background, SCOT+ unifies single-cell optimal transport into the most digestible framework since the topic’s inception for the single-cell domain.

SCOT+ will enable biologists to make additional progress towards inferring gene regulatory networks as well as producing more robust cross-modal genomic models. Specifically, these downstream analyses require co-assay datasets that are available through the use of SCOT+, but are otherwise hard to generate experimentally. Additionally, biologists will able to better infer cell trajectories during development or cellular evolution across species.

In the future, we are looking into the theoretical implications of unbalancing the AGW formulation as we have in this package. Additionally, we are continuing to expand SCOT+ to additional real-world scenarios to aid users in tuning SCOT+ for their specific needs.

## Data and Code Availability

All of these results, including thorough tutorials on the theory and application of each formulation as well as practical examples on real-world data, are available in downloadable Jupyter notebooks on our website, hosted at https://scotplus.github.io/. The website additionally contains documentation that will help users use this tool on their own data.

The PBMC dataset can be found at https://www.10xgenomics.com/datasets/pbmc-from-a-healthy-donor-granulocytes-removed-through-cell-sorting-3-k-1-standard-2-0-0. The CITE-seq dataset can be found at https://www.nature.com/articles/nmeth.4380.

## Acknowledgments

This work is supported by the National Institutes of Health (NIH) Award 1R35HG011939-01 and Brown University’s Undergraduate Teaching and Research Award (UTRA). It is also supported by the Brown Center for Computation and Visualization.

