## Supplementary Materials for "SCOT+: A Comprehensive Software Suite for Single-Cell alignment Using Optimal Transport"

### SCOT+ Supplementary Materials

#### 1 Optimization Procedure

##### 1.1 Overview

To solve the minimization problems given by the formulations in the methods section, SCOT+ utilizes a general solution which solves the unbalanced AGW minimization problem. This solution follows from the observation that we can solve an unbalanced OT problem for $P$ when $P'$ and $Q$ are frozen by reorganizing our original loss function [1]:

$$\begin{aligned} L &= \alpha GW(D_x, D_y) + (1-\alpha)COOT(X,Y)\big) \\ &= \langle C, P\rangle + \varepsilon_{P',Q} KL(P|\mu_{sx}\otimes\mu_{sy}) \\ &\quad + \rho_{1P',Q} KL(P_{\#1}|\mu_{sx}) + \rho_{2P',Q} KL(P_{\#2}|\mu_{sy}) \end{aligned}$$

As we can see, this reorganization of our loss yields a standard unbalanced OT problem. However, note that we introduced some new variables ($\varepsilon_{P',Q}, \rho_{iP',Q}, C$) to package each of these unbalanced OT parameters more cleanly. These variables have the following definitions, starting with the regularization:

$$\varepsilon_{P',Q} = \alpha\varepsilon_{gw} m(P') + (1-\alpha)\varepsilon_{coot} m(Q)$$

In this case, we see that the regularization of the UOT problem we found balances the underlying regularization of the GW and COOT problems. We see a similar pattern for the relaxation parameters:

$$\rho_{iP',Q} = \alpha\rho_i^{gw} M(P') + (1-\alpha)\rho_i^{coot} M(Q) \text{ for } i = 1,2$$

We can see that, like the regularization parameter, the relaxation parameter of this UOT problem balances the relaxation of each subproblem according to $\alpha$. Finally, holding $P'$ and $Q$ constant allow us to find the cost matrix:

$$\begin{aligned} C = \alpha\Big[ & [D_x^{\odot 2} P'_{\#1} \oplus D_y^{\odot 2} P'_{\#2} + 2D_x P' D_y^T] \\ & + \varepsilon_{gw}\langle \log(\frac{P'}{\mu_{sx}\otimes\mu_{sy}}), P'\rangle \\ & + \rho_1^{gw}\langle \log(\frac{P'_{\#1}}{\mu_{sx}}), P'_{\#1}\rangle + \rho_2^{gw}\langle \log(\frac{P'_{\#2}}{\mu_{sy}}), P'_{\#2}\rangle\Big] \\ & + (1-\alpha)\Big[[X^{\odot 2} Q_{\#1} \oplus Y^{\odot 2} Q_{\#2} + 2XQY^T] \\ & + \varepsilon_{coot}\langle \log(\frac{Q}{\mu_{fx}\otimes\mu_{fy}}), Q\rangle \\ & + \rho_1^{coot}\langle \log(\frac{Q_{\#1}}{\mu_{fx}}), Q_{\#1}\rangle + \rho_2^{coot}\langle \log(\frac{Q_{\#2}}{\mu_{fy}}), Q_{\#2}\rangle\Big] + \beta_s D_s \end{aligned}$$

Before diving into the details of finding $C$ below, we can get some intuition for where each term comes from. First, notice that there is a GW component associated with $P'$ multiplied by $\alpha$ and a COOT component associated with $Q$ multiplied by $(1-\alpha)$. Within each of these wider components, we can see that we factor out $P'$ and $Q$ from the main inner product term, the regularization term, and the relaxation term of the AGW problem. This forms three terms per subproblem, amounting to the first six terms in the above definition. Finally, similar to other OT problems, supervision is easy to package into this cost matrix, which then gives the seventh and final term.

The above analysis is an example of how we can transform an unbalanced AGW problem into a standard unbalanced OT problem by holding two of the three coupling matrices constant. More generally, when freezing all but one coupling matrix (not necessarily just $P$) as above in our more complicated OT formulations, we recover a standard unbalanced optimal transport problem which we can solve via Sinkhorn's algorithm.

By repeatedly iterating over solving the resulting UOT problems for each of $P$, $P'$, and $Q$, we recover the set of coupling matrices that minimizes the unbalanced AGW loss. This iterative procedure is reminiscent of the block coordinate descent algorithm [1]. Finally, as shown in 1.7 below, each SCOT+ formulation can now be recovered by using appropriate values for $\alpha$ and $\rho$.

#### 1.2 Formalizing the SCOT+ Objective Function

In order to compute the cost-minimizing coupling matrices for each formulation in SCOT+, we approximate the arguments $P, Q, P'$ that minimize the following equation:

$$\begin{aligned} L = \alpha\Big[&\langle |D_x - D_y|^2, P \otimes P'\rangle + \beta_s\langle D_s, P\rangle + \beta_s\langle D_s, P'\rangle + \rho_1^{gw} KL(P_{\#1} \otimes P'_{\#1} | \mu_{sx} \otimes \mu_{sx}) \\ &+ \rho_2^{gw} KL(P_{\#2} \otimes P'_{\#2} | \mu_{sy} \otimes \mu_{sy}) + \varepsilon_{gw} KL(P \otimes P' | \mu_{sx} \otimes \mu_{sy} \otimes \mu_{sx} \otimes \mu_{sy})\Big] \\ &+ (1-\alpha)\Big[\langle |X - Y|^2, P \otimes Q\rangle + \beta_s\langle D_s, P\rangle + \beta_f\langle D_f, Q\rangle + \rho_1^{coot} KL(P_{\#1} \otimes Q_{\#1} | \mu_{sx} \otimes \mu_{fx}) \\ &+ \rho_2^{coot} KL(P_{\#2} \otimes Q_{\#2} | \mu_{sy} \otimes \mu_{fy}) + \varepsilon_{coot} KL(P \otimes Q | \mu_{sx} \otimes \mu_{sy} \otimes \mu_{fx} \otimes \mu_{fy})\Big] \end{aligned}$$

Note here the following notation concerns:

1. $D_x$ and $D_y$ are distance matrices computed from $X$ and $Y$. We generally compute these by constructing a $k$nn graph and using Dijkstra's algorithm, which we provide in our utils module (`compute_graph_distances`).

2. $\beta_s$ and $D_s$ are the supervision hyperparameters described in our methods.

3. The $x$ in $\rho_x$ refers to which domain we regularize/relax, while the $y$ in $\rho^y$ refers to which type of OT formulation the term modifies.

4. The $i$ in $\mu_{ij}$ refers to whether the related measure supports a marginal distribution in the sample or feature domain. The $j$ refers to which domain the measure supports ($x$ aligns with 1 domain-wise for say, $\rho_1$ and $\mu_x$; $y$ aligns with domain 2).

5. $\otimes$ is the Kronecker product.

6. $C_{\#x}$ indicates the marginal distribution of $C$ along dimension $x$. So, for example, $P_{\#2}$ is the marginal distribution of the sample coupling matrix along the domain 2 axis; i.e., $P_{\#2}$ describes the total mass transported by each sample in domain 2. Mathematically, $P_{\#2} = \sum_i P_{ij}$ for discrete joint distributions.

Interpreting this equation makes the most sense element-wise; the large terms of which we are taking a convex combination are UGW and UCOOT loss terms respectively. Within each, we have the main UOT cost, as well as supervision, relaxation, and regularization costs. Note that we could also have looked at

$$
\begin{aligned}
L = \alpha \Big[ &\langle |D_x - D_y|^2, P \otimes P' \rangle + \beta_s \langle D_s, P \rangle + \beta_s \langle D_s, P' \rangle + \rho_1^{gw} KL(P_{\#1} \otimes P'_{\#1} | \mu_{sx} \otimes \mu_{sx}) \\
&+ \rho_2^{gw} KL(P_{\#2} \otimes P'_{\#2} | \mu_{sy} \otimes \mu_{sy}) + \varepsilon_{gw} KL(P | \mu_{sx} \otimes \mu_{sy}) + \varepsilon_{gw} KL(P' | \mu_{sx} \otimes \mu_{sy}) \Big] \\
&+ (1-\alpha) \Big[ \langle |X - Y|^2, P \otimes Q \rangle + \beta_s \langle D_s, P \rangle + \beta_f \langle D_f, Q \rangle + \rho_1^{coot} KL(P_{\#1} \otimes Q_{\#1} | \mu_{sx} \otimes \mu_{fx}) \\
&+ \rho_2^{coot} KL(P_{\#2} \otimes Q_{\#2} | \mu_{sy} \otimes \mu_{fy}) + \varepsilon_{gw} KL(P | \mu_{sx} \otimes \mu_{sy}) + \varepsilon_{coot} KL(Q | \mu_{fx} \otimes \mu_{fy}) \Big]
\end{aligned}
$$

for independent entropic regularization, which introduces more hyperparameters (unless you set those that modify matrices that should be similar equal, as above) but gives slightly more control over coupling matrix density. Since the calculations with joint entropic regularization are slightly more involved and easily extended to independent regularization, we'll focus on the joint entropic regularization case.

In order to approximate the cost-minimizing $P, Q, P'$, we utilize a form of block coordinate descent (BCD) where we hold two coupling matrices constant and solve the resulting UOT problem with respect to the third matrix. We can examine how this makes the minimization problem solvable by reorganizing the loss function under each circumstance.

#### 1.3 Solving for $P$

To solve for a given one of the three coupling matrices, we need to solve a minimization problem that holds the other two matrices constant. As such, once we treat two of the three coupling matrices as constant, we can get rid of all terms that do not contain the one "free" matrix, as these terms are constant with respect to our minimization. The $\rightarrow$ symbols indicate dropping a number of constants related to the two frozen matrices.

$$
\begin{aligned}
L = \alpha\Big[&\langle |D_x - D_y|^2, P\otimes P'\rangle + \beta_s\langle D_s, P\rangle + \beta_s\langle D_s, P'\rangle + \rho_1^{gw}KL(P_{\#1}\otimes P'_{\#1}|\mu_{sx}\otimes\mu_{sx}) \\
&+ \rho_2^{gw}KL(P_{\#2}\otimes P'_{\#2}|\mu_{sy}\otimes\mu_{sy}) + \varepsilon_{gw}KL(P\otimes P'|\mu_{sx}\otimes\mu_{sy}\otimes\mu_{sx}\otimes\mu_{sy})\Big] \\
&+ (1-\alpha)\Big[\langle |X-Y|^2, P\otimes Q\rangle + \beta_s\langle D_s, P\rangle + \beta_f\langle D_f, Q\rangle + \rho_1^{coot}KL(P_{\#1}\otimes Q_{\#1}|\mu_{sx}\otimes\mu_{fx}) \\
&+ \rho_2^{coot}KL(P_{\#2}\otimes Q_{\#2}|\mu_{sy}\otimes\mu_{fy}) + \varepsilon_{coot}KL(P\otimes Q|\mu_{sx}\otimes\mu_{sy}\otimes\mu_{fx}\otimes\mu_{fy})\Big] \\
\rightarrow &\langle\alpha\Big[[D_x^{\odot 2}P'_{\#1}\oplus D_y^{\odot 2}P'_{\#2} + 2D_xP'D_y^T] + \varepsilon_{gw}\langle\log(\frac{P'}{\mu_{sx}\otimes\mu_{sy}}), P'\rangle\Big] \\
&+ (1-\alpha)\Big[[X^{\odot 2}Q_{\#1}\oplus Y^{\odot 2}Q_{\#2} + 2XQY^T] + \varepsilon_{coot}\langle\log(\frac{Q}{\mu_{fx}\otimes\mu_{fy}}), Q\rangle\Big] + \beta_sD_s, P\rangle \\
&+ \alpha\varepsilon_{gw}m(P')KL(P|\mu_{sx}\otimes\mu_{sy}) + (1-\alpha)\varepsilon_{coot}m(Q)KL(P|\mu_{sx}\otimes\mu_{sy}) \\
&+ \alpha\Big(\rho_1^{gw}KL(P_{\#1}\otimes P'_{\#1}|\mu_{sx}\otimes\mu_{sx}) + \rho_2^{gw}KL(P_{\#2}\otimes P'_{\#2}|\mu_{sy}\otimes\mu_{sy})\Big) \\
&+ (1-\alpha)\Big(\rho_1^{coot}KL(P_{\#1}\otimes Q_{\#1}|\mu_{sx}\otimes\mu_{fx}) + \rho_2^{coot}KL(P_{\#2}\otimes Q_{\#2}|\mu_{sy}\otimes\mu_{fy})\Big) \\
\rightarrow &\langle\alpha\Big[[D_x^{\odot 2}P'_{\#1}\oplus D_y^{\odot 2}P'_{\#2} + 2D_xP'D_y^T] + \varepsilon_{gw}\langle\log(\frac{P'}{\mu_{sx}\otimes\mu_{sy}}), P'\rangle\Big] \\
&+ (1-\alpha)\Big[[X^{\odot 2}Q_{\#1}\oplus Y^{\odot 2}Q_{\#2} + 2XQY^T] + \varepsilon_{coot}\langle\log(\frac{Q}{\mu_{fx}\otimes\mu_{fy}}), Q\rangle\Big] + \beta_sD_s, P\rangle \\
&+ \alpha\varepsilon_{gw}m(P')KL(P|\mu_{sx}\otimes\mu_{sy}) + (1-\alpha)\varepsilon_{coot}m(Q)KL(P|\mu_{sx}\otimes\mu_{sy}) \\
&+ \alpha\Big(\rho_1^{gw}\Big[m(P')KL(P_{\#1}|\mu_{sx}) + \langle\langle\log(\frac{P'_{\#1}}{\mu_{sx}}), P'_{\#1}\rangle, P\rangle\Big] + \rho_2^{gw}\Big[m(P')KL(P_{\#2}|\mu_{sy}) + \langle\langle\log(\frac{P'_{\#2}}{\mu_{sy}}), P'_{\#2}\rangle, P\rangle\Big]\Big) \\
&+ (1-\alpha)\Big(\rho_1^{coot}\Big[m(Q)KL(P_{\#1}|\mu_{sx}) + \langle\langle\log(\frac{Q_{\#1}}{\mu_{fx}}), Q_{\#1}\rangle, P\rangle\Big] + \rho_2^{coot}\Big[m(Q)KL(P_{\#2}|\mu_{sy}) + \langle\langle\log(\frac{Q_{\#2}}{\mu_{fy}}), Q_{\#2}\rangle, P\rangle\Big]\Big) \\
= &\langle C, P\rangle + \Big(\alpha\varepsilon_{gw}m(P') + (1-\alpha)\varepsilon_{coot}m(Q)\Big)KL(P|\mu_{sx}\otimes\mu_{sy}) \\
&+ \Big(\alpha\rho_1^{gw}M(P') + (1-\alpha)\rho_1^{coot}M(Q)\Big)KL(P_{\#1}|\mu_{sx}) + \Big(\alpha\rho_2^{gw}M(P') + (1-\alpha)\rho_2^{coot}M(Q)\Big)KL(P_{\#2}|\mu_{sy})
\end{aligned}
$$

Where

$$
\begin{aligned}
C = \alpha\Big[&[D_x^{\odot 2}P'_{\#1}\oplus D_y^{\odot 2}P'_{\#2} + 2D_xP'D_y^T] + \varepsilon_{gw}\langle\log(\frac{P'}{\mu_{sx}\otimes\mu_{sy}}), P'\rangle \\
&+ \rho_1^{gw}\langle\log(\frac{P'_{\#1}}{\mu_{sx}}), P'_{\#1}\rangle + \rho_2^{gw}\langle\log(\frac{P'_{\#2}}{\mu_{sy}}), P'_{\#2}\rangle\Big] \\
&+ (1-\alpha)\Big[[X^{\odot 2}Q_{\#1}\oplus Y^{\odot 2}Q_{\#2} + 2XQY^T] + \varepsilon_{coot}\langle\log(\frac{Q}{\mu_{fx}\otimes\mu_{fy}}), Q\rangle \\
&+ \rho_1^{coot}\langle\log(\frac{Q_{\#1}}{\mu_{fx}}), Q_{\#1}\rangle + \rho_2^{coot}\langle\log(\frac{Q_{\#2}}{\mu_{fy}}), Q_{\#2}\rangle\Big] + \beta_sD_s
\end{aligned}
$$

Note that the first and second steps follow by the observation (as in Proposition 4 of [2]) that

$$\begin{aligned}
\text{reg./relax. cost} &= KL(P_{\#1} \otimes X_{\#1} | \mu_{sx} \otimes \nu_1) \\
&= \sum_{i,j} \Big( P_{\#1_i} X_{\#1_j} \log \frac{P_{\#1_i} X_{\#1_j}}{\mu_{sx_i} \nu_{1j}} \Big) - \sum_{i,j} P_{\#1_i} X_{\#1_j} + \sum_{i,j} \mu_{sx_i} \nu_{1j} \\
&= \sum_{i,j} \Big( P_{\#1_i} X_{\#1_j} \big( \log \frac{P_{\#1_i}}{\mu_{sx_i}} + \log \frac{X_{\#1_j}}{\nu_{1j}} \big) \Big) - m(P)m(X) + m(\mu_{sx})m(\nu_1) \\
&= \sum_{i,j} \Big( P_{\#1_i} X_{\#1_j} \log \frac{X_{\#1_j}}{\nu_{1j}} \Big) + \sum_{i,j} \Big( P_{\#1_i} X_{\#1_j} \log \frac{P_{\#1_i}}{\mu_{sx_i}} \Big) \\
&\quad - m(P)m(X) + m(\mu_{sx})m(\nu_1) + \big( m(P)m(\nu_1) - m(P)m(\nu_1) \big) \\
&= m(P) KL(X_{\#1} | \nu_1) + \sum_{i,j} \Big( P_{\#1_i} X_{\#1_j} \log \frac{P_{\#1_i}}{\mu_{sx_i}} \Big) \\
&\quad + m(\mu_{sx})m(\nu_1) - m(P)m(\nu_1) \\
&= m(P) KL(X_{\#1} | \nu_1) + \sum_{i,j} \Big( P_{\#1_i} X_{\#1_j} \log \frac{P_{\#1_i}}{\mu_{sx_i}} \Big) + m(\mu_{sx})m(\nu_1) - m(P)m(\nu_1) \\
&\quad + \big( m(P)m(X) - m(P)m(X) \big) + \big( m(X)m(\mu_{sx}) - m(X)m(\mu_{sx}) \big) \\
&= m(P) KL(X_{\#1} | \nu_1) + m(X) KL(P_{\#1} | \mu_{sx}) \\
&\quad + m(\mu_{sx})m(\nu_1) - m(P)m(\nu_1) + m(P)m(X) - m(X)m(\mu_{sx}) \\
&= m(P) KL(X_{\#1} | \nu_1) + m(X) KL(P_{\#1} | \mu_{sx}) + \big[ m(P) - m(\mu_{sx}) \big] \big[ m(X) - m(\nu) \big] \\
&= \int \Big( \int \log \big( \frac{dX_{\#1}}{d\nu_1} \big) dX_{\#1} \Big) dP + m(X) KL(P_{\#1} | \mu_{sx}) - m(\mu_{sx}) \big[ m(X) - m(\nu_1) \big] \\
&= \langle \Big( \int \log \big( \frac{dX_{\#1}}{d\nu_1} \big) dX_{\#1} \Big), P \rangle + m(X) KL(P_{\#1} | \mu_{sx}) + \text{ constant w.r.t. } P \\
&\to \underbrace{\langle \langle \log \big( \frac{X_{\#1}}{\nu_1} \big), X_{\#1} \rangle, P \rangle}_{\text{will go into } C} + \underbrace{m(X) KL(P_{\#1} | \mu_{sx})}_{\text{will be part of core UOT}}
\end{aligned}$$

for $X = Q, P'$ with respective supports $\nu_1, \nu_2$ and that this also extends to the case where we have terms of the form $KL(P \otimes X | \mu_{sx} \otimes \mu_{sy} \otimes \nu_1 \otimes \nu_2)$:

$$\begin{aligned}
\text{reg./relax. cost} &= KL(P \otimes X | (\mu_{sx} \otimes \mu_{sy}) \otimes (\nu_1 \otimes \nu_2)) \\
&= m(P) KL(X | \nu_1 \otimes \nu_2) + m(X) KL(P | \mu_{sx} \otimes \mu_{sy}) \\
&\quad + \big[ m(P) - m(\mu_{sx} \otimes \mu_{sy}) \big] \big[ m(X) - m(\nu_1 \otimes \nu_2) \big] \\
&= \int \Big( \int \log \big( \frac{dX}{d\nu_1 \otimes \nu_2} \big) dX \Big) dP + m(X) KL(P | \mu_{sx} \otimes \mu_{sy}) \\
&\quad - m(\mu_{sx} \otimes \mu_{sy}) \big[ m(X) - m(\nu_1 \otimes \nu_2) \big] \\
&= \langle \Big( \int \log \big( \frac{dX}{d\nu_1 \otimes \nu_2} \big) dX \Big), P \rangle + m(X) KL(P | \mu_{sx} \otimes \mu_{sy}) + \text{ constant w.r.t. } P \\
&\to \underbrace{\langle \langle \log \big( \frac{X}{\nu_1 \otimes \nu_2} \big), X \rangle, P \rangle}_{\text{will go into } C} + \underbrace{m(X) KL(P | \mu_{sx} \otimes \mu_{sy})}_{\text{will be part of core UOT}}
\end{aligned}$$

Looking at the final result of this calculation, we have a UOT problem with a cost matrix $C$ that we can compute. We can also compute each scalar multiplier of the regularization and relaxation terms, such that we are essentially solving a problem of form

$$\langle C, P\rangle + \varepsilon^* KL(P|\mu_{sx} \otimes \mu_{sy}) + \rho_1^* KL(P_{\#1}|\mu_{sx}) + \rho_2^* KL(P_{\#2}|\mu_{sy})$$

Which can be solved via Sinkhorn iteration, as per [1] and [3]. We will call one set of Sinkhorn (scaling) iterations to solve for $P$ a call to the function `IsolateP`. We can do the same manipulation, even more simply, for $P'$ and $Q$.

#### 1.4 Solving for P'

$$\begin{aligned}
L = \alpha\Big[&\langle |D_x - D_y|^2, P \otimes P'\rangle + \beta_s\langle D_s, P\rangle + \beta_s\langle D_s, P'\rangle + \rho_1^{gw} KL(P_{\#1} \otimes P'_{\#1}|\mu_{sx} \otimes \mu_{sx}) \\
&+ \rho_2^{gw} KL(P_{\#2} \otimes P'_{\#2}|\mu_{sy} \otimes \mu_{sy}) + \varepsilon_{gw} KL(P \otimes P'|\mu_{sx} \otimes \mu_{sy} \otimes \mu_{sx} \otimes \mu_{sy})\Big] \\
&+ (1-\alpha)\Big[\langle |X - Y|^2, P \otimes Q\rangle + \beta_s\langle D_s, P\rangle + \beta_f\langle D_f, Q\rangle + \rho_1^{coot} KL(P_{\#1} \otimes Q_{\#1}|\mu_{sx} \otimes \mu_{fx}) \\
&+ \rho_2^{coot} KL(P_{\#2} \otimes Q_{\#2}|\mu_{sy} \otimes \mu_{fy}) + \varepsilon_{coot} KL(P \otimes Q|\mu_{sx} \otimes \mu_{sy} \otimes \mu_{fx} \otimes \mu_{fy})\Big] \\
\to \alpha\Big[&\langle |D_x - D_y|^2, P \otimes P'\rangle + \rho_1^{gw} KL(P_{\#1} \otimes P'_{\#1}|\mu_{sx} \otimes \mu_{sx}) \\
&+ \rho_2^{gw} KL(P_{\#2} \otimes P'_{\#2}|\mu_{sy} \otimes \mu_{sy}) + \beta_s\langle D_s P'\rangle + \varepsilon_{gw} KL(P \otimes P'|\mu_{sx} \otimes \mu_{sy} \otimes \mu_{sx} \otimes \mu_{sy})\Big] \\
\to \alpha\Big[&\langle [D_x^{\odot 2} P_{\#1} \oplus D_y^{\odot 2} P_{\#2} + 2D_x P D_y^T] + \varepsilon_{gw}\langle \log(\frac{P}{\mu_{sx} \otimes \mu_{sy}}), P\rangle + \beta_s D_s, P'\rangle + \varepsilon_{gw} m(P) KL(P'|\mu_{sx} \otimes \mu_{sy}) \\
&+ \rho_1^{gw} KL(P_{\#1} \otimes P'_{\#1}|\mu_{sx} \otimes \mu_{sx}) + \rho_2^{gw} KL(P_{\#2} \otimes P'_{\#2}|\mu_{sy} \otimes \mu_{sy})\Big] \\
= \alpha\Big[&\langle [D_x^{\odot 2} P_{\#1} \oplus D_y^{\odot 2} P_{\#2} + 2D_x P D_y^T] + \varepsilon_{gw}\langle \log(\frac{P}{\mu_{sx} \otimes \mu_{sy}}) + \beta_s D_s, P\rangle, P'\rangle + \varepsilon_{gw} m(P) KL(P'|\mu_{sx} \otimes \mu_{sy}) \\
&+ \rho_1^{gw} m(P) KL(P'_{\#1}|\mu_{sx}) + \rho_2^{gw} m(P) KL(P'_{\#2}|\mu_{sy}) + \rho_1^{gw}\langle\langle \log(\frac{P_{\#1}}{\mu_{sx}}), P_{\#1}\rangle, P'\rangle + \rho_2^{gw}\langle\langle \log(\frac{P_{\#2}}{\mu_{sy}}), P_{\#2}\rangle, P'\rangle\Big] \\
= \langle C, P'\rangle &+ \alpha\varepsilon_{gw} m(P) KL(P'|\mu_{sx} \otimes \mu_{sy}) + \alpha\rho_1^{gw} m(P) KL(P'_{\#1}|\mu_{sx}) + \alpha\rho_2^{gw} m(P) KL(P'_{\#2}|\mu_{sy})
\end{aligned}$$

Where

$$\begin{aligned}
C = \alpha\Big[&[D_x^{\odot 2} P_{\#1} \oplus D_y^{\odot 2} P_{\#2} + 2D_x P D_y^T] + \varepsilon_{gw}\langle \log(\frac{P}{\mu_{sx} \otimes \mu_{sy}}), P\rangle + \\
&+ \rho_1^{gw}\langle \log(\frac{P_{\#1}}{\mu_{sx}}), P_{\#1}\rangle + \rho_2^{gw}\langle \log(\frac{P_{\#2}}{\mu_{sy}}), P_{\#2}\rangle + \beta_s D_s\Big]
\end{aligned}$$

We will call one set of Sinkhorn iterations to solve for $P'$ a call to the function `IsolateP'`.

#### 1.5 Solving for Q

$$\begin{aligned}
L &= \alpha\Big[\langle |D_x - D_y|^2, P\otimes P'\rangle + \beta_s\langle D_s, P\rangle + \beta_s\langle D_s, P'\rangle + \rho_1^{gw} KL(P_{\#1}\otimes P'_{\#1}|\mu_{sx}\otimes\mu_{sx}) \\
&\quad + \rho_2^{gw} KL(P_{\#2}\otimes P'_{\#2}|\mu_{sy}\otimes\mu_{sy}) + \varepsilon_{gw} KL(P\otimes P'|\mu_{sx}\otimes\mu_{sy}\otimes\mu_{sx}\otimes\mu_{sy})\Big] \\
&\quad + (1-\alpha)\Big[\langle |X-Y|^2, P\otimes Q\rangle + \beta_s\langle D_s, P\rangle + \beta_f\langle D_f, Q\rangle + \rho_1^{coot} KL(P_{\#1}\otimes Q_{\#1}|\mu_{sx}\otimes\mu_{fx}) \\
&\quad + \rho_2^{coot} KL(P_{\#2}\otimes Q_{\#2}|\mu_{sy}\otimes\mu_{fy}) + \varepsilon_{coot} KL(P\otimes Q|\mu_{sx}\otimes\mu_{sy}\otimes\mu_{fx}\otimes\mu_{fy})\Big] \\
&\to (1-\alpha)\Big[\langle |X-Y|^2, P\otimes Q\rangle + \beta_s\langle D_s, P\rangle + \beta_f\langle D_f, Q\rangle + \rho_1^{coot} KL(P_{\#1}\otimes Q_{\#1}|\mu_{sx}\otimes\mu_{fx}) \\
&\quad + \rho_2^{coot} KL(P_{\#2}\otimes Q_{\#2}|\mu_{sy}\otimes\mu_{fy}) + \varepsilon_{coot} KL(P\otimes Q|\mu_{sx}\otimes\mu_{sy}\otimes\mu_{fx}\otimes\mu_{fy})\Big] \\
&= (1-\alpha)\Big[\langle [X^{T^{\odot 2}}P_{\#1}\oplus Y^{T^{\odot 2}}P_{\#2} + 2X^TPY] + \varepsilon_{coot}\langle \log(\frac{P}{\mu_{sx}\otimes\mu_{sy}}) + \beta_f D_f, P\rangle, Q\rangle + \varepsilon_{coot} m(P) KL(Q|\mu_{fx}\otimes\mu_{fy}) \\
&\quad + \rho_1^{coot} KL(P_{\#1}\otimes Q_{\#1}|\mu_{sx}\otimes\mu_{fx}) + \rho_2^{coot} KL(P_{\#2}\otimes Q_{\#2}|\mu_{sy}\otimes\mu_{fy})\Big] \\
&= (1-\alpha)\Big[\langle [X^{T^{\odot 2}}P_{\#1}\oplus Y^{T^{\odot 2}}P_{\#2} + 2X^TPY] + \varepsilon_{coot}\langle \log(\frac{P}{\mu_{sx}\otimes\mu_{sy}}), P\rangle + \beta_f D_f, Q\rangle + \varepsilon_{coot} m(P) KL(Q|\mu_{fx}\otimes\mu_{fy}) \\
&\quad + \rho_1^{coot} m(P) KL(Q_{\#1}|\mu_{fx}) + \rho_2^{coot} m(P) KL(Q_{\#2}|\mu_{fy}) + \rho_1^{coot}\langle\langle \log(\frac{P_{\#1}}{\mu_{sx}}), P_{\#1}\rangle, Q\rangle + \rho_2^{coot}\langle\langle \log(\frac{P_{\#2}}{\mu_{sy}}), P_{\#2}\rangle, Q\rangle\Big] \\
&= \langle C, Q\rangle + (1-\alpha)\varepsilon_{coot} m(P) KL(Q|\mu_{fx}\otimes\mu_{fy}) + (1-\alpha)\rho_1^{coot} m(P) KL(Q_{\#1}|\mu_{fx}) + (1-\alpha)\rho_2^{coot} m(P) KL(Q_{\#2}|\mu_{fy})
\end{aligned}$$

Where

$$\begin{aligned}
C = (1-\alpha)\Big[&[X^{T^{\odot 2}}P_{\#1}\oplus Y^{T^{\odot 2}}P_{\#2} + 2X^TPY] + \varepsilon_{coot}\langle \log(\frac{P}{\mu_{sx}\otimes\mu_{sy}}), P\rangle \\
&+ \rho_1^{coot}\langle \log(\frac{P_{\#1}}{\mu_{sx}}), P_{\#1}\rangle + \rho_2^{coot}\langle \log(\frac{P_{\#2}}{\mu_{sy}}), P_{\#2}\rangle + \beta_f D_f\Big]
\end{aligned}$$

We will call one set of Sinkhorn iterations to solve for $Q$ a call to the function `IsolateQ`.

Note that, rather than explicitly optimize for $P'$ as well, we could instead have alternated freezing $Q$ and solving the resulting FUGW problem with freezing $P, P'$ and solving the resulting UOT problem for $Q$.

##### 1.6 General Procedure

Note that in the below, the following inputs are tuples:

1. $\beta = (\beta_s, \beta_f)$
2. $D = (D_s, D_f)$
3. $\rho^{gw} = (\rho_1^{gw}, \rho_2^{gw})$
4. $\rho^{coot} = (\rho_1^{coot}, \rho_2^{coot})$
5. $\mu_x = (\mu_{sx}, \mu_{sy})$
6. $\mu_y = (\mu_{fx}, \mu_{fy})$

Additionally, we rescale as per [1].

**Algorithm 1** General SCOT+ Algorithm

**procedure** SOLVE($X, Y, D_x, D_y, \alpha, \beta, D, \varepsilon_{gw}, \varepsilon_{coot}, \rho^{gw}, \rho^{coot}, \mu_s, \mu_f$)
    **for** $1 \leq i \leq n_{bcd}$ **do**
        **for** $1 \leq j \leq n_{gw}$ **do**
            $P' \leftarrow$ `IsolateP'`$(P, D_x, D_y, \alpha, \beta, D, \varepsilon_{gw}, \rho^{gw}, \mu_s)$
            $P' \leftarrow \sqrt{\frac{m(P)}{m(P')}} P'$
            $P \leftarrow$ `IsolateP`$(P', Q, X, Y, D_x, D_y, \alpha, \beta, D, \varepsilon_{gw}, \varepsilon_{coot}, \rho^{gw}, \rho^{coot}, \mu_s, \mu_f)$
            $P \leftarrow \sqrt{\frac{m(P')}{m(P)}} P$
        $Q \leftarrow$ `IsolateQ`$(P, X, Y, \alpha, \beta, D, \varepsilon_{coot}, \rho^{coot}, \mu_s, \mu_f)$
        $Q \leftarrow \sqrt{\frac{m(P)}{m(Q)}} Q$
        $P \leftarrow$ `IsolateP`$(P', Q, X, Y, D_x, D_y, \alpha, \beta, D, \varepsilon_{gw}, \varepsilon_{coot}, \rho^{gw}, \rho^{coot}, \mu_s, \mu_f)$
        $P \leftarrow \sqrt{\frac{m(Q)}{m(P)}} P$
    **return** $P, P', Q$

##### 1.7 Isolating Formulations

Our current package supports direct use of the general framework presented in supplementary materials section 1.1 via calls to `uagw`, but additionally allows for calls to older optimal transport formulations useful for multi-omic data alignment.

#### 1.7.1 GW

In particular, we can partial out the GW formulation [4] by solving the UAGW problem with $\alpha = 1, \rho \to \infty$ for all $\rho$:

$$\begin{aligned}
L = \alpha\Big[&\langle |D_x - D_y|^2, P \otimes P'\rangle + \beta_s\langle D_s, P\rangle + \beta_s\langle D_s, P'\rangle + \rho_1^{gw} KL(P_{\#1} \otimes P'_{\#1} | \mu_{sx} \otimes \mu_{sx}) \\
&+ \rho_2^{gw} KL(P_{\#2} \otimes P'_{\#2} | \mu_{sy} \otimes \mu_{sy}) + \varepsilon_{gw} KL(P \otimes P' | \mu_{sx} \otimes \mu_{sy} \otimes \mu_{sx} \otimes \mu_{sy})\Big] \\
&+ (1-\alpha)\Big[\langle |X - Y|^2, P \otimes Q\rangle + \beta_s\langle D_s, P\rangle + \beta_f\langle D_f, Q\rangle + \rho_1^{coot} KL(P_{\#1} \otimes Q_{\#1} | \mu_{sx} \otimes \mu_{fx}) \\
&+ \rho_2^{coot} KL(P_{\#2} \otimes Q_{\#2} | \mu_{sy} \otimes \mu_{fy}) + \varepsilon_{coot} KL(P \otimes Q | \mu_{sx} \otimes \mu_{sy} \otimes \mu_{fx} \otimes \mu_{fy})\Big] \\
= &\langle |D_x - D_y|^2, P \otimes P'\rangle + \beta_s\langle D_s, P\rangle + \beta_s\langle D_s, P'\rangle + \rho_1^{gw} KL(P_{\#1} \otimes P'_{\#1} | \mu_{sx} \otimes \mu_{sx}) \\
&+ \rho_2^{gw} KL(P_{\#2} \otimes P'_{\#2} | \mu_{sy} \otimes \mu_{sy}) + \varepsilon_{gw} KL(P \otimes P' | \mu_{sx} \otimes \mu_{sy} \otimes \mu_{sx} \otimes \mu_{sy})
\end{aligned}$$

Since we allow $\rho_1^{gw}$, $\rho_2^{gw} \to \infty$, we enforce that $P_{\#1} = \mu_{sx}, P_{\#2} = \mu_{sy}$ and the equivalent for $P'$, otherwise $L \to \infty$. In particular, $P, P'$ must both be in the set $\Pi_s = \{X | \sum_i X_{ij} = \mu_{sx}, \sum_j X_{ij} = \mu_s y\}$. So, to minimize $L$ with respect to $P$ and $P'$, we find $P$ and $P'$ with the fixed (balanced) marginal supports specified by $\mu_{sx}$ and $\mu_{sy}$ to minimize the following

$$L_{gw} = \langle |D_x - D_y|^2, P \otimes P'\rangle + \beta_s\langle D_s, P\rangle + \beta_s\langle D_s, P'\rangle + \varepsilon_{gw} KL(P \otimes P' | \mu_{sx} \otimes \mu_{sy} \otimes \mu_{sx} \otimes \mu_{sy})$$

Which is exactly the GW problem if we set $\beta_s = 0$, and otherwise the fused Gromov-Wasserstein problem (FGW). Technically, this is a restricted version of FGW that forces what would be $\beta'_s$ equal to $\beta_s$ and $D'_s$ equal to $D_s$, but since we want $P' \to P$, we fix these equalities for simplicity.

##### 1.7.2 UGW

If we then let our $\rho$ variables vary but fix $\alpha = 1$, we recover

$$\begin{aligned}
L = &\langle |D_x - D_y|^2, P \otimes P'\rangle + \beta_s\langle D_s, P\rangle + \beta_s\langle D_s, P'\rangle + \rho_1^{gw} KL(P_{\#1} \otimes P'_{\#1} | \mu_{sx} \otimes \mu_{sx}) \\
&+ \rho_2^{gw} KL(P_{\#2} \otimes P'_{\#2} | \mu_{sy} \otimes \mu_{sy}) + \varepsilon_{gw} KL(P \otimes P' | \mu_{sx} \otimes \mu_{sy} \otimes \mu_{sx} \otimes \mu_{sy})
\end{aligned}$$

from our GW derivation, which is exactly the UGW problem from [5] if we set $\beta_s = 0$, and otherwise the fused unbalanced Gromov-Wasserstein problem (FUGW).

##### 1.7.3 COOT

In the parallel case where we fix $\alpha = 0$ and allow all $\rho \to \infty$, we recover the COOT problem [6].

$$\begin{aligned}
L = \alpha\Big[&\langle |D_x - D_y|^2, P \otimes P'\rangle + \beta_s\langle D_s, P\rangle + \beta_s\langle D_s, P'\rangle + \rho_1^{gw} KL(P_{\#1} \otimes P'_{\#1} | \mu_{sx} \otimes \mu_{sx}) \\
&+ \rho_2^{gw} KL(P_{\#2} \otimes P'_{\#2} | \mu_{sy} \otimes \mu_{sy}) + \varepsilon_{gw} KL(P \otimes P' | \mu_{sx} \otimes \mu_{sy} \otimes \mu_{sx} \otimes \mu_{sy})\Big] \\
&+ (1-\alpha)\Big[\langle |X - Y|^2, P \otimes Q\rangle + \beta_s\langle D_s, P\rangle + \beta_f\langle D_f, Q\rangle + \rho_1^{coot} KL(P_{\#1} \otimes Q_{\#1} | \mu_{sx} \otimes \mu_{fx}) \\
&+ \rho_2^{coot} KL(P_{\#2} \otimes Q_{\#2} | \mu_{sy} \otimes \mu_{fy}) + \varepsilon_{coot} KL(P \otimes Q | \mu_{sx} \otimes \mu_{sy} \otimes \mu_{fx} \otimes \mu_{fy})\Big] \\
= &\langle |X - Y|^2, P \otimes Q\rangle + \beta_s\langle D_s, P\rangle + \beta_f\langle D_f, Q\rangle + \rho_1^{coot} KL(P_{\#1} \otimes Q_{\#1} | \mu_{sx} \otimes \mu_{fx}) \\
&+ \rho_2^{coot} KL(P_{\#2} \otimes Q_{\#2} | \mu_{sy} \otimes \mu_{fy}) + \varepsilon_{coot} KL(P \otimes Q | \mu_{sx} \otimes \mu_{sy} \otimes \mu_{fx} \otimes \mu_{fy})
\end{aligned}$$

Since our $\rho$ variables approach infinity, any valid minimizer to $L$ must have $P \in \Pi_s$ and $Q \in \Pi_f$, where $\Pi_f = \{X | \sum_i X_{ij} = \mu_{fx}, \sum_j X_{ij} = \mu_{fy})\}$. Additionally, within this constraint, $(P, Q)$ minimize

$$L_{coot} = \langle |X - Y|^2, P \otimes Q \rangle + \beta_s \langle D_s, P \rangle + \beta_f \langle D_f, Q \rangle + \varepsilon_{coot} KL(P \otimes Q | \mu_{sx} \otimes \mu_{sy} \otimes \mu_{fx} \otimes \mu_{fy})$$

such that $P$ and $Q$ solve the COOT problem when $\beta_s, \beta_f = 0$ and the fused unbalanced co-optimal transport (FCOOT) problem otherwise.

##### 1.7.4 UCOOT

Parallel to UGW, if we let our $\rho$ variables vary but fix $\alpha = 0$, we recover

$$\begin{aligned} L = \langle |X - Y|^2, P \otimes Q \rangle + \beta_s \langle D_s, P \rangle + \beta_f \langle D_f, Q \rangle + \rho_1^{coot} KL(P_{\#1} \otimes Q_{\#1} | \mu_{sx} \otimes \mu_{fx}) \\ + \rho_2^{coot} KL(P_{\#2} \otimes Q_{\#2} | \mu_{sy} \otimes \mu_{fy}) + \varepsilon_{coot} KL(P \otimes Q | \mu_{sx} \otimes \mu_{sy} \otimes \mu_{fx} \otimes \mu_{fy}) \end{aligned}$$

from our COOT derivation, which is exactly the UCOOT problem [1] if we set $\beta_s, \beta_f = 0$, and otherwise the fused unbalanced co-optimal transport problem (FUCOOT).

##### 1.7.5 AGW

Finally, by allowing $\rho \to \infty$ for all $\rho$ in our generalized equation, we recover the augmented Gromov-Wasserstein (AGW) problem from [7]. In particular, when we fully enforce all of the marginal constraints rather than relaxing, we drop all $\rho$ terms just as in the GW and COOT derivations, which leaves

$$\begin{aligned} L_{agw} = \alpha \Big[ \langle |D_x - D_y|^2, P \otimes P' \rangle + \beta_s \langle D_s, P \rangle + \beta_s \langle D_s, P' \rangle + \varepsilon_{gw} KL(P \otimes P' | \mu_{sx} \otimes \mu_{sy} \otimes \mu_{sx} \otimes \mu_{sy}) \Big] \\ + (1 - \alpha) \Big[ \langle |X - Y|^2, P \otimes Q \rangle + \beta_s \langle D_s, P \rangle + \beta_f \langle D_f, Q \rangle + \varepsilon_{coot} KL(P \otimes Q | \mu_{sx} \otimes \mu_{sy} \otimes \mu_{fx} \otimes \mu_{fy}) \Big] \end{aligned}$$

Meaning that when we allow $\rho \to \infty$ in our generalized solver approach, we choose $P, P' \in \Pi_s$ and $Q \in \Pi_f$ to minimize $L_{agw}$. In other words, we recover solutions to the AGW problem. Each of these sections has shown that from our most general SCOT+ solver, we can recover solutions to each of the GW, UGW, COOT, UCOOT, and AGW problems for given tabular datasets $X, Y$.

#### 1.8 Software

SCOT+ makes accessing solutions to each of the formulations above easier than ever. Now available on the Python package index, `scotplus` has an easy interface that users can interact with once their data is loaded into Python in numpy [8], pandas [9], or torch [10]. In order to align two domains, users can follow the steps below:

1. Import `solvers.SinkhornSolver`, our current solver for each of these formulations.

2. Instantiate a `SinkhornSolver` object, which requires a number of optimization-related hyperparamaters, including number of iterations, convergence tolerance, and frequency of convergence evaluation.

3. Decide on the right formulation for the domains to be aligned. For example, if we were working with data that had disproportionate cell type representation and unclear feature relationships, we might choose UGW.

4. If the formulation chosen includes any GW terms, compute the intra-domain distance matrices for each domain. We provide a built-in option that we have found works best, which can be imported via `utils.alignment.compute_graph_distances` and run directly on each domain. This particular method computes a nearest-neighbor graph within each domain and produces an intra-domain distance matrix.

5. Using the previously instantiated `SinkhornSolver`, call any of the built-in alignment methods, which include `gw, ugw, coot, ucoot`, `agw`, and `uagw`, on your data with their default parameters. Each of these methods takes in some combination of count matrices and intra-domain distance matrices in order to compute a sample and feature coupling matrix.

6. Tune the hyperparameters of the alignment method chosen. In particular, users might need to tune the $\varepsilon$, $\rho$, and $\alpha$ parameters described above, as well as some optimization hyperparameters that are less crucial to the results. Running SCOT+ formulations with a small $\varepsilon$ to $\rho$ ratio can take up to an hour on a local CPU, but such solutions are generally too sparse. In our experience, an optimal set of hyperparameters will take at most 30 minutes to run on a local machine.

7. Pass the anchor domain count matrix and the sampling coupling matrix returned by the alignment method into `utils.alignment.get_barycentre`. This method returns the samples of the source domain projected into the feature space of the anchor domain.

8. In examples with some ground-truth information, score the final alignment produced by running `utils.alignment.FOSCTTM` or `utils.alignment.LTA`. Each of these scoring schemes is described below and requires some amount of prior knowledge to execute.

#### 2 Hyperparameter Tuning

See below for a list of hyperparameters used to generate the figures in the the body of the main paper. Note that all examples in the body of the paper were fully unsupervised, i.e. $\beta_s = \beta_f = 0$.

*Cell-cell alignment on balanced datasets*
$\boldsymbol{\varepsilon = 0.002}$
*Cell-cell alignment on unbalanced datasets*
$\boldsymbol{\varepsilon = 0.00127, \rho_1 = \rho_2 = 0.01}$
*Feature-based alignment on balanced datasets*
$\boldsymbol{\alpha = 0.1, \varepsilon_{gw} = \varepsilon_{coot} = 0.00166}$
*Feature-based alignment on unbalanced datasets*
$\boldsymbol{\alpha = 0.1, \varepsilon_{gw} = 0.002, \varepsilon_{coot} = 0.001, \rho_1^{gw} = \rho_2^{gw} = \infty, \rho_1^{coot} = \rho_2^{coot} = 0.005}$

More generally, we recommend tuning by first running a grid search over $\varepsilon_{gw}$ and $\varepsilon_{coot}$ until you reach a reasonable level of density in your sample and feature coupling matrices. From here, we recommend running another grid search over $\rho_{gw}$ and $\rho_{coot}$, where in each case we set $\rho_x^1 = \rho_x^2$, depending on whether you think your populations are unbalanced in some way (i.e., disproportionate cell type representation, disproportionate feature contributions/relevance). Finally, if you are tuning with AGW, we recommend following the same steps, but additionally searching over a narrow range for $\alpha$, generally further towards COOT (i.e., $0.1 \leq \alpha \leq 0.5$) in order to provide further resolution on feature relationships. Sometimes, if you are using our `compute_graph_distances` function, you may also want to tune $k$; we had it set to 110 for both PBMC applications and 100 for both CITE-seq applications. See our website as listed below for more information.

As an example, in our UGW application to the PBMC dataset, we searched over $\rho_{gw}^1 = \rho_{gw}^2 \in \{0.01, 0.1, 1\}$ jointly with $\varepsilon_{gw} \in \{10^{-3}, 10^{-2.9}, ...10^{-1}\}$.

### 3 Practical Alignment

The optimization procedure explored in the first section of the supplementary materials returns three matrices: $P, P'$, and $Q$. However, at first glance, these matrices do not clearly give an alignment across datasets. However, we can project domain 1 (i.e., has samples along the $i$ axis of $P_{ij}$) onto domain 2 (samples along $j$ axis of $P_{ij}$) using $P$. Let $X \in \mathbb{R}^{n_x \times d_x}$ be the tabular data for domain 1, and $Y \in \mathbb{R}^{n_y \times d_y}$ be the tabular data for domain 2. Additionally, let $\hat{Y} \in \mathbb{R}^{n_x \times d_y}$ be the target data, i.e. approximate values for the samples in $X$ for the features of $Y$. Then, as in [1],

$$\hat{Y}_i = \underset{y \in \mathbb{R}^{d_y}}{\text{argmin}} \big( \sum_{j=1}^{n_y} P_{ij} c(y, y_j) \big)$$

Which has a closed form solution when we chose $c$ to be squared Euclidean distance,

$$\hat{Y}_i = \sum_{j=1}^{n_y} \frac{P_{ij}}{P_{\#1_i}} Y_j$$

As a result, we can recover $\hat{Y}$ directly from $P$. From here, we can continue to analyze the recovered alignment by concatenating $X$ and $\hat{Y}$ to have an approximate tabular dataset with $n_x$ samples and $d_x + d_y$ features. We choose $P$ instead of $P'$ given that $P$ contributes to minimizing both UGW and UCOOT losses. This function is available in our `utils` module via the function `get_barycentre`.

### 4 More Information

In order to get a sense for how SCOT+ works in more practical scenarios, please visit our website, which we currently host here ((`https://scotplus.github.io/`). The website has downloadable tutorials and documentation for the tool. Additionally, SCOT+ will soon be available on PyPI via the command `pip install scotplus`.
